# The influence of reproductive mode on resource competition and diversity patterns in Ediacaran early animal communities

**DOI:** 10.1101/2025.01.08.632049

**Authors:** Emily G. Mitchell, Andrea Manica

## Abstract

The appearance of the oldest known animals during the late Ediacaran (∼574 million years ago [Ma])^1–4^ was followed by a phase of little change^5^. This period, which lasted ∼14 million years, ended with a burst of rapid diversification known as the Ediacaran “Second Wave”. The reasons for these diversity patterns are poorly understood. Here we investigate how reproductive mode mediated community dynamics, and in turn macroevolutionary change, in the Ediacaran. We show that widespread reproduction via stolon (namely via filaments connecting clones) in the first animals limited intra-specific competition among neighbours, leading to inter-specific competition acting at smaller-spatial scales than intra-specific competition, a phenomenon called heteromyopia^6^. Heteromyopia enables co-existence of sub-optimal competitors because the dispersal limitation of the dominant species means that they do not inhabit all the optimal habitat, so that lesser competitors can still exist within the same community, operating under reduced selection pressure. We explored the consequences of this dispersal limitation on community diversity using Approximate Bayesian Computation to estimate the posterior distributions of dispersal with a spatially explicit model fitted to the three Ediacaran assemblages and showed that the change from stoloniferous to sexual reproduction that coincided with the Second Wave could explain the sudden increase in alpha diversity observed in the fossil record. We conclude that widespread asexual reproduction via stolon likely constrained early animal evolution, limiting diversification until the onset of mobility and widespread sexual reproduction^7–9^.

## Main Text

One of the most dramatic events in the evolutionary history of Earth is the appearance of animals in the fossil record during the Ediacaran time period (635-539 Ma) after billions of years of microbial life^1–4^. The earliest widespread animal communities are represented as Ediacaran macrofossils of Avalonia (574 – 560 Ma)^2,4,10^, consisting of in-situ, sessile benthic organisms^11–13^. This in-situ preservation of sessile benthic communities enables the use of spatial analyses to resolve reproductive modes^14^, habitat associations^15^ and resource competition^15–17^ within Avalonian communities. These spatial analyses have shown that the driving forces behind resource competition are not well understood. Whilst modern benthic communities are highly competitive, driven by competition for space and/or food (e.g. Ref. ^18^), such competition is rare and weak in Avalonian communities^15–17^. Competition for water column resources, such as food, are unlikely because there are no correlations between the presence of inter-specific competition and specimen height, demonstrating that these taxa do not avoid competing for resources from the same part of the water column^15^ Taken together, these results suggest that Avalonian systems are compatible with neutral models^17^. However, in the Avalon there are instances of inter-specific competition occurring at shorter spatial scales than intra-specific competition^17^, a pattern known as heteromyopia^6^ which suggests other key processes may be involved^6,19^. Importantly, heteromyopic dynamics cannot be distinguished from neutral dynamics based on most ecological metrics, such as species area distributions^20^, or spatial point process analyses, the main tools used to test the fit of neutral models which have shown predominately neutral dynamics for Avalonian communities^17^. In extant communities, heteromyopia, (while rare^19^) can be caused by dispersal limitation due to connected stoloniferous clusters, pathogens, predators, allopathic death or strong niche segregation^6^. Only by modelling explicitly the spatial scales of interactions, something that has not yet been done for Ediacaran communities, can we directly quantify heteromyopic dynamics, and thus move beyond the niche-neutral paradigm of previous work to explore whether a third process, reproductive mode, mediates the community dynamics.

Stoloniferous reproduction has been directly observed in five Avalonian taxa^21^, and its occurrence was first inferred based on the size-structured spatial arrangement of specimens^14^, supported by numerical and computational fluid dynamics^22^. Stoloniferous reproduction has the capacity to reduce intra-specific competition as follows: if stoloniferous clones remain connected by stolon throughout their lives, they would share nutrients (regardless of feeding mode, or height of feeding) between these clones, thus acting as a unit and negating the need for nutrient competition between individuals. If one clone is in a low resource area, it can gain nutrients via stolon from clones in higher resource areas. As such, competition will not occur within connected stoloniferous networks, it would only occur between clonal colonies since each connected clonal colony is acting as a unit. Therefore, stoloniferous reproduction could be a mechanism for the relatively low levels of intra-specific competition and heteromyopia that is found within Avalonian communities^15,16^ which has the potential to lead to low alpha diversity (species richness) levels^23^.

In this study we test the importance of intra-specific processes, specifically stoloniferous reproduction, to the presence and strength of intra-specific resource competition (i.e. quantify the strength of heteromyopic dynamics) and use mechanistic models to investigate how these processes may have impacted the biodiversity of early animal communities of the Ediacaran (∼574-539 Ma).

### Identification and quantification of stoloniferous reproduction and resource competition

Stoloniferous reproduction restricts the direction and distance offspring can travel from their parents as in animals they remain attached via these stolon to their parents. This attachment results in isotropic clusters, that is clusters which do not have any directionality, i.e. are circular rather than more ovate, and also results in small clusters sizes because the offspring cannot be transported long distances by currents (as per Mitchell et al. 2015^14^ and Delahooke et al., 2024^22^). On the other hand, where benthic organisms disperse pelagically transported by currents (either via pelagic larvae, or pelagic buds or fragments), we expect clusters to be elongated, following the direction of the current (Extended data Fig. 1). In order for a signal of stoloniferous reproduction to be found, the population should also be relatively mature, so the relative age of the population may also impact the ability to find such reproductive modes^14^.

We developed a metric to quantify cluster directionality by measuring the interquartile range of the orientations of all specimens relative to each other, with isotropy indicated by small values, and anisotropy by high values. Stoloniferous connection between individuals will limit the size of dispersal clusters, since water-borne propagules travel further than connected ones^24–27^, leading to smaller cluster sizes. Because such reproductive cluster sizes depend on the height of the parent, here we consider normalised clusters (i.e. we divide the cluster size by the maximum specimen height of population) rather than absolute sizes. The third factor which would influence the prevalence of stoloniferous reproduction is the relative maturity of the population. Well-established populations will have had more time to reproduce, so are likely to have higher percentages of the second/third generations. If new generations result from stoloniferous reproduction, then we would expect to see stronger effects of stoloniferous reproduction for more mature communities. Therefore, as a proxy for relative age of a population, we used the percentage of the population that occupies the smallest size-class, with the larger the percentage, the younger the population.

Crucial to determining the relationship between reproductive mode and community dynamics is determining the strength of competition within these communities. While identifying the most likely process underlying a spatial pattern is not straightforward^28–32^, competition within sessile communities can be identified as the most likely underlying process through spatial segregation which is not linked to underlying habitat segregation^33,34^.The presence of resource competition has previously been detected within Avalonian communities using spatial point process analyses (SPPA)^14,16,17^, identifying four cases of inter-specific competition within seven communities. However, these studies focussed on detecting competition, rather estimating its strength^14,17,35^. Here we use SPPA to generate a metric of competition strength and use it to investigate its drivers. SPPA can be used to quantify the level of spatial segregation that is not caused by habitat association^33,34,36^: specifically, we use the distance measure Pair Correlation Functions (PCF) to quantify how the density of points (here fossil specimens) changes across the mapped area (here bedding planes)^37^ of 20 populations across 8 bedding planes from the Avalon of Newfoundland, Canada and Charnwood Forest, UK, using pre-existing data^14,17,35,37^. PCF = 1 represents complete spatial randomness, with values below one corresponding to spatial segregation, above 1 to spatial aggregation. The strength of segregation, i.e. resource competition, is indicated by the smallest PCF value, i.e. the minimum value of the PCF, and the spatial scale over which resource competition occurs is the segregation up until (or over) the point where the observed data crosses PCF = 1^37^.

### Relationship of reproductive mode to resource competition

We started by applying our new directionality measure to 20 taxa populations (data from Ref. 17). There was a range of cluster directionality (Fig. 1), with H14 *Fractofusus andersoni* showing the lowest levels of isotropy and Ostrich Feathers on LMP showing the highest levels of anisotrophy (Fig 1). The *Fractofusus andersoni* result is consistent with previous work that used spatial analyses to infer stoloniferous reproduction^14^, as well as physical evidence of filaments connected to specimens^21^. In contrast, the Bristy Cove *Fractofusus* showed high levels of directionality, likely due to the relative immaturity of the community (as inferred by the small body sizes and large percentage in the smallest size class, Table 1), suggesting that the population had not had not enough time to reproduce. Ostrich Feathers have been found with connected filaments^21^, but the relatively small body-sizes of the specimens on Lower Mistaken Point Surface (6^th^ smallest maximum height, Table 1) suggest it is likely to also be an immature community, with a lack of widespread second-generation reproduction. The Bed B *Primocandelabrum boyntoni* was an outlier for non-segregation populations with high isotopy (Table 1, Fig. 1). Bed B is thought to be dominated by a secondary-succession^38^, whereby the largest specimens are survivors from an earlier death-event so this combination of multiple populations may explain this mixed signal. The breadth of directionality results (Table 1) demonstrates that stoloniferous reproduction was occurring in varying amounts across multiple bedding planes, enabling further analyses to be able to test the consequences of such reproductive mode.

**Figure 1:**
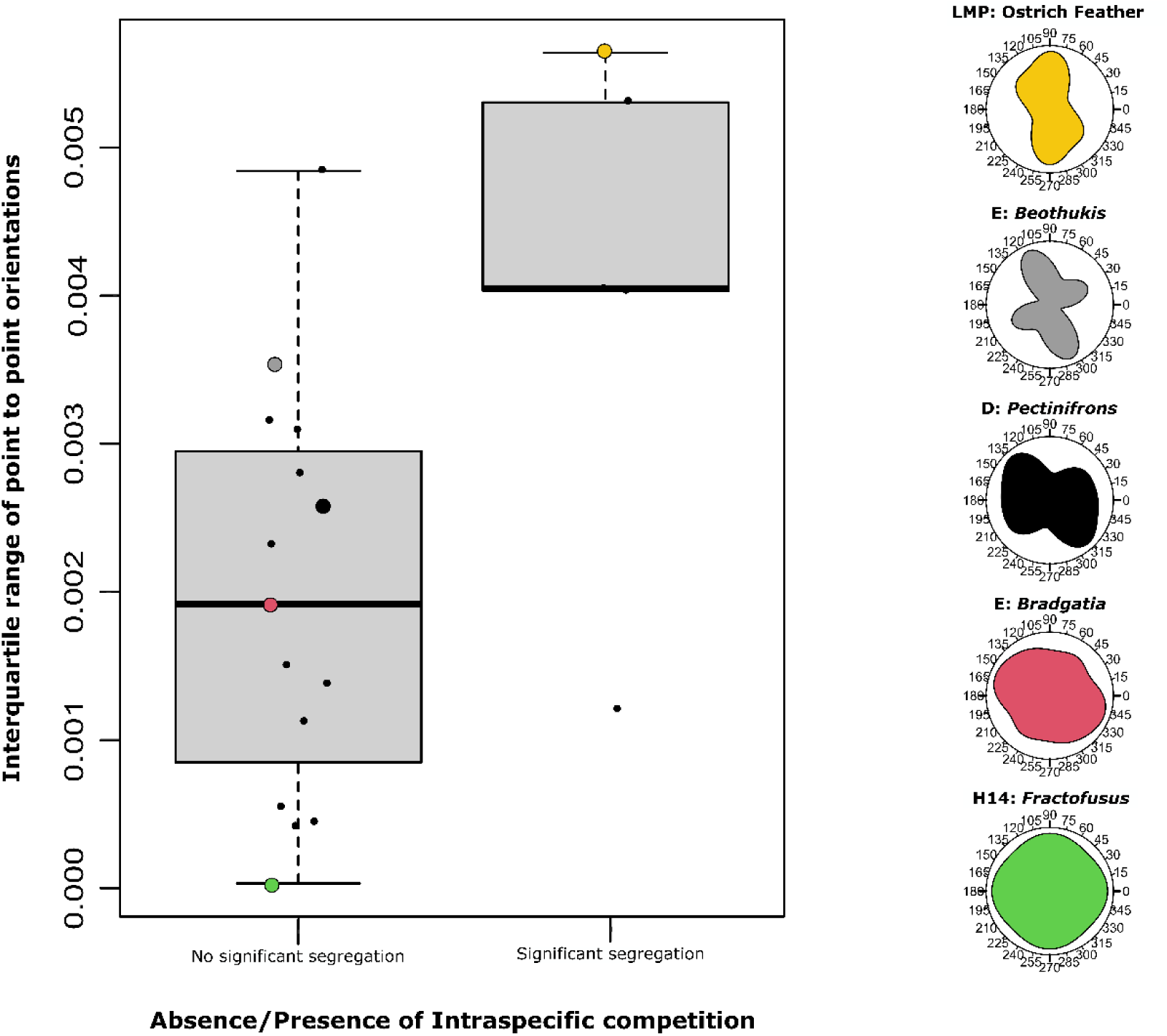
Boxplot of interquartile range of point to point angles grouped depending on whether they exhibit significant segregation. Lower IQR indicates isotropy, and higher IQR anisotropy. Example rose diagrams are given on the right, with the corresponding IQR given on the boxplot in the same colour.

We quantified the strength and spatial scale of intra-specific competition, using the approach developed in Mitchell et al. 2019^14^. There were 16 populations which were non-random, of which five taxa populations had excursions under the Monte Carlo simulation envelopes, so were considered to exhibit intra-specific competition (Fig. 2). G surface *Bradgatia* exhibited the strongest segregation (PCF_min_ = 0.384) and Bed B *Charnia* the weakest of the significant segregations (PCF_min_ = 0.692), with a mean PCF_min_ = 0.604 for the populations which exhibited intra-specific spatial segregation (Table 1). There were eight cases of significant inter-specific segregation, of which only one had inter-specific segregation on a larger spatial scales than intra-specific segregation (Fig. 2). There were three inter-specific distributions where the spatial scale of the minimum PCF was smaller than for one species, and three where it was smaller than both species (Fig. 2). There was one instance with *Plumeropriscum plumosa - Charniodiscus spinous* where the inter-specific spatial scale of *Charniodiscus spinous* was comparable to that of the inter-specific segregation. The largest heteromyopic difference of spatial scales was with *Fractofusus* and *Charniodiscus procerus* on E surface, where the inter-specific segregation occurred 1.21m before the intra-specific segregation of *Charniodiscus procerus* and 2.85m than *Fractofusus* (Fig. 2). These results demonstrate the heteromyopic relationships across these communities (Fig. 2).

**Figure 2:**
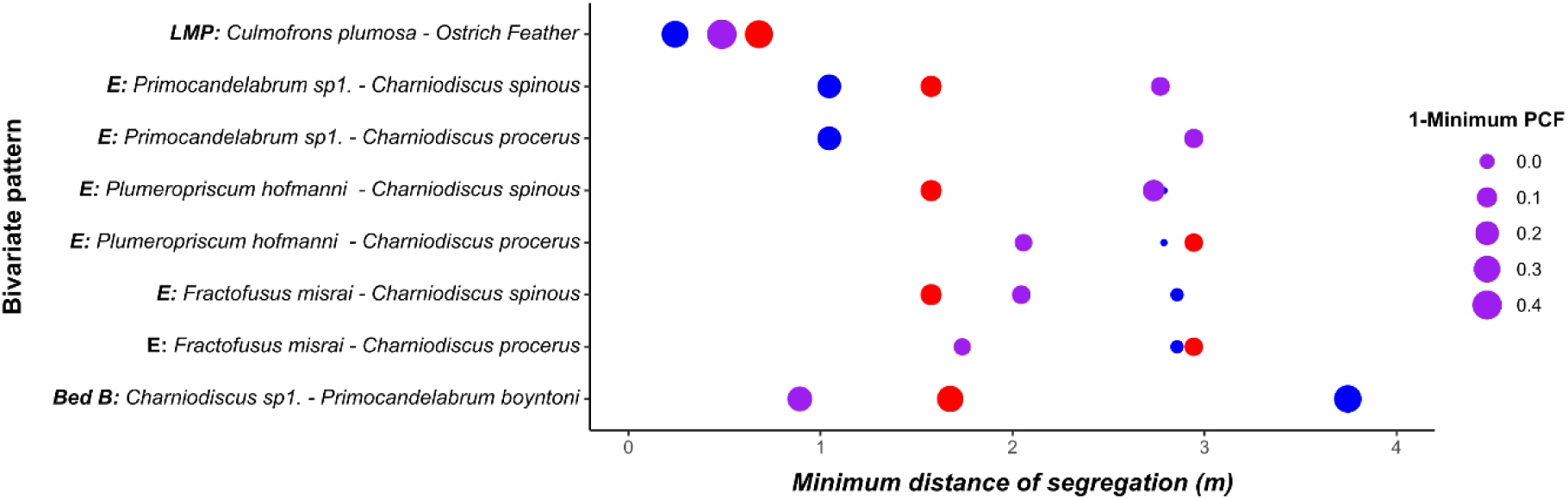
Plot of the Minimum distance of segregation for each non-random bivariate spatial pattern. Red points are the first species of the pattern, Blue points depict the second species pattern and purple the values for the bivariate (interspecific) pattern. The size of the points corresponds to the relative strength of the segregation, given by 1-PCF_min_. See Extended data figure 2 for the PCFs plots from which these data were extracted. Heteromyopic relationships were present if the interspecific estimate (purple point) was found at smaller minimum distance than either of the intra-specific once (blue/red).

The quantification of factors associated with stoloniferous reproduction (Fig. 1), coupled with the quantification of the strength of intra-specific segregation (Fig. 2, Extended data Fig. 2), enables us to systemically test whether there was an association between stoloniferous reproduction (as indicated by our directionality metric) and the presence of intra-specific segregation with, through a Mann-Whitney test to test for differences of the directionality metric between the presence/absence groups for intra-specific segregation. We found a significant association between the presence/absence of this intra-specific segregation with cluster directionality (W = 63, N_1_ = 15, N_2_ = 5, *p* = 0.0254) (Fig. 1), suggesting that stoloniferous reproduction inhibits the presence of intra-specific competition. We found that the strength of intraspecific segregation was best predicted by considering cluster directionality, normalised cluster size and percentage of the population in smallest size class (F_3,16_ = 7.722, *p =* 0.0021, R^2^=0.515; Extended data Table 3). Most of the explanatory power came from cluster directionality and normalised cluster size (F_2,17_ = 9.937, *p*=0.0013, R^2^=0.488, Fig. 3. ΔAIC = 0.4181). These regressions suggest that the strength of intra-specific competition is reduced within stoloniferous populations.

**Figure 3:**
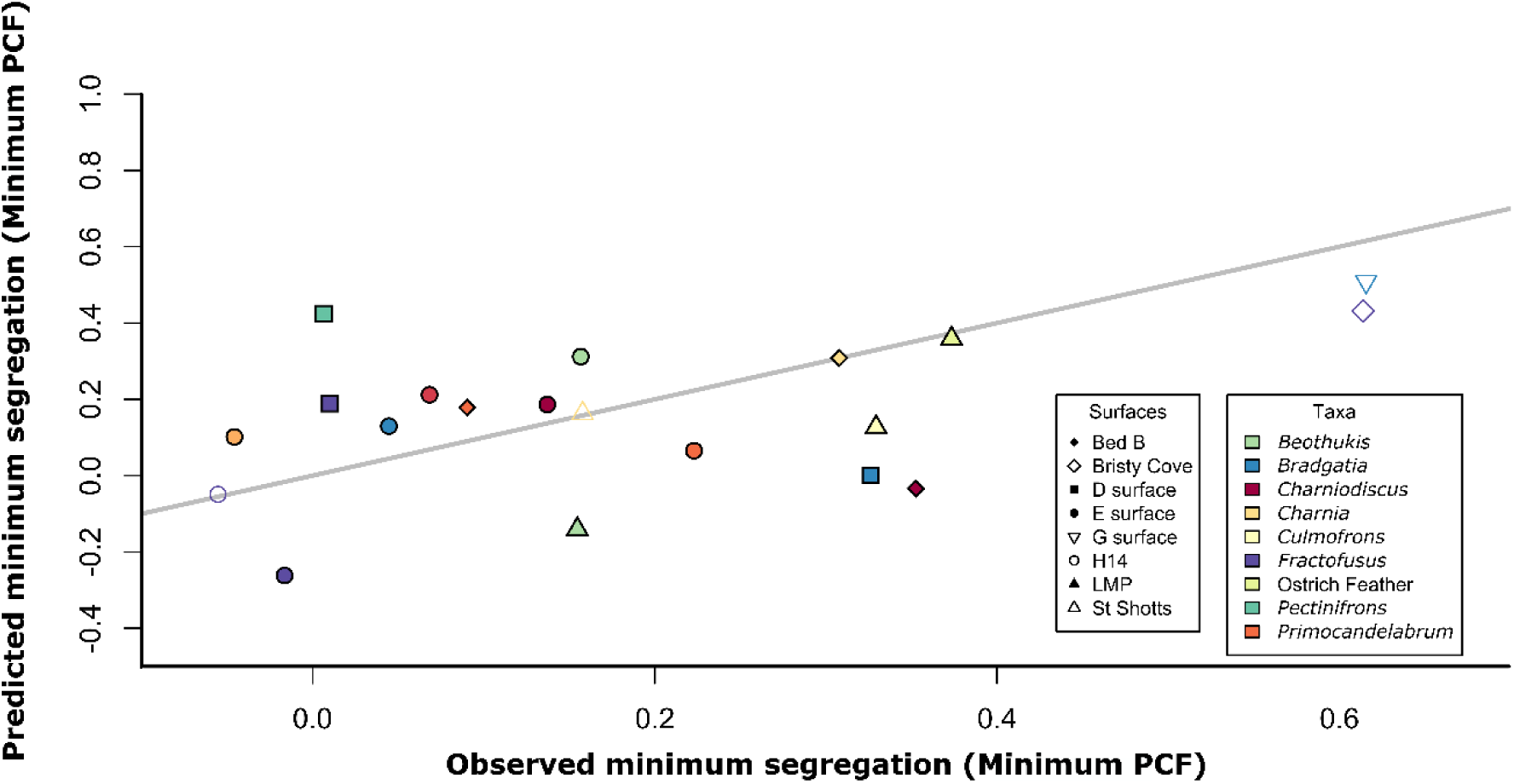
Relationship between observed and predicted competition as indicated by the minimum PCF. The x axis is the observed minimum PCFs and the simulated minimum is given on the y axis using the best-fit regression model with the two variables of interquartile range of the directionality and normalised clusters sizes (*p*=0.0013, R^2^=0.488). Line of best fit is shown in grey.

Of the eight communities, only four had sufficiently abundant multi-taxa populations to enable bivariate analyses. Of these four, three communities had non-random spatial distributions (Fig. 2). Within these three communities, there are four instances of bivariate distributions where the spatial scale of inter-specific competition occurs at smaller spatial scales than intra-specific competition for both taxa (Fig. 2) with two further instances where inter-specific competition was detected at smaller spatial scales than intra-specific competition for one ttaxon indicating heteromyopia for one of the two taxa (Fig. 2). Therefore, there are six instances of heteromyopic spatial patterns, i.e. all three communities which had non-random bivariate spatial patterns exhibited heteromyopia. In terms of the five processes that can lead to heteromyopia, namely stoloniferous clusters, pathogens, predators, allopathic death or strong niche segregation^6^, there is no evidence of macro-predation until the terminal Ediacaran, making predation unlikely. Niche influences on Avalonian taxa are rare and weak^15,17^, and there is no evidence of Avalonian niche segregation^39^, so niche segregation is also unlikely. The highly repetitive nature of Avalonian spatial patterns across potentially large space and time intervals^17^ also makes niche processes less likely. Allelopathy and pathogens would be expected to have stronger effects at the smallest spatial scales, because the chemicals (or pathogens) have the highest density around the individual, so would lead to the largest segregations at small spatial scales, which a decrease as spatial scale increases. Such strong, small-scale segregation is not observed in any of the eight communities (Fig. 2, Extended data Fig. 2). Because there is a significant association between factors associated with stoloniferous reproduction and resource competition (indicated by spatial segregation), we suggest that connected stoloniferous networks drive the heteromyopic patterns found.

### Consequences of heteromyopia on Ediacaran diversification patterns

Our data is a good representation of Avalonian life because the communities we studied include 76% of documented species from the Avalon^40^, covering 572 – 560 Ma. In this study, we analysed all the abundant populations, which accounted for 46% of documented species with a further 30% of species found within the mapped areas but not in sufficient abundances for these analyses (n > 30) and the remaining 26% of taxa considered rare (< 10 documented specimens)^40,41^. As such, our data is a good reflection of documented early animal life. Within our studied communities, where resource competition occurs (indicated by spatial segregation), heteromyopia occurs in six out of eight cases, including between the most abundant species populations, so is likely a dominant force behind community dynamics.

Heteromyopia changes the dynamics of communities in terms of selection pressures on good and bad competitors^6^. In non-dispersal limited communities, the strongest competitor will take over all the optimal habitat, leading to competition-colonization trade-offs, strong selection pressures, and the best competitors leaving little room for sub-optimal competitors to establish themselves^43^. However, where dispersal is strongly limited, such as for connected stoloniferous networks, populations occupy a mixture of good and bad resources, as long as at least some of the connected stoloniferous network is in the good habitat^44,45^. Unlike dispersal unlimited taxa, which are able to spread widely and colonize all the optimal resources, dispersal limited taxa cannot spread so easily, so often optimal resources are still available to sub-optimal competitors^46^. This dynamic creates a mechanism of co-existence whereby weaker competitors co-exist alongside much stronger competitor taxa^46^, and so are under reduced selection pressure^6,47^. Our results suggest that in these Avalonian communities, stoloniferous reproduction leads to heteromyopia, which dominates the community dynamics. Yet, the majority of taxa (70%) are not abundant, so do not form a significant proportion of the communities, and are not subject to these heteromyopic processes, and instead enjoy an easier existence, with reduced selection pressure. This heteromyopic dynamic is consistent with the accumulation mode of community development^48^. In such communities, instead of communities developing systematically in response to deterministic set of inter-specific interactions, the composition of the communities do not change through time, and instead are dependent on the initial colonisers, whose reproductive mode shapes how the communities mature.

The lack of selection pressure for the majority of Avalonian taxa due to heteromyopia may explain the relative low rates of Avalonian diversification, with the majority of taxa not under strong evolutionary pressure, but the dominant taxa exerting a heteromyopic dynamic, thus enabling co-existence, but without a strong driver for natural selection. This co-existence dynamic would change either through a change in reproductive strategy, from stoloniferous to water-borne reproduction (cf. Ref. 49) and/or through the breakage of the stoloniferous connections that could come through the advent of mobility animals mining in, on and under the microbial mats that covered the Ediacaran substrate^50,51^. The stolon-breakage would lead to intra-specific competition at smaller spatial-scales^52^, thus increasing competition and selection pressure for both the previously heteromyopic organisms as well as the weaker competitors that had previously been shielded by the dispersal limited dominant taxa. Thus, increase in Ediacaran diversity around 550 Ma^7^, has the potential be explained from the increase of dispersal distances^53^ and subsequent changes of selection pressure that comes with a transition from wide-spread stoloniferous reproduction, to systems with waterborne propagation^49,53^, and mobile organisms^50^ breaking stolon, and thus the prior co-existence mechanism.

To investigate the impact of dispersal limitation on alpha diversity we constructed a mechanistic model of the Avalon, White Sea and Nama assemblages cf. Ref. 54. These three assemblages represent the taxonomic grouping of Ediacaran macrofossil taxa, with each assemblage occupying largely different temporal periods and different environmental settings^40,55^ and often are used when discussing Ediacaran evolutionary patterns to distinguish different evolutionary phrases. These three assemblages show a dramatic increase in alpha diversity between the Avalon and White Sea assemblages, with a more limited decrease in the Nama^56^. In order to see the extent to which heteromyopia can explain these trends we created a mechanistic model to simulate alpha diversity. Our model was a spatially explicit lottery metacommunity model^54^, parameterised by the extent of immigration between communities, spatial autocorrelation (i.e. the extent of association with habitat heterogeneities), distance of dispersal and niche overlap. Given the observed values of alpha diversity for the Avalon, White Sea and Nama assemblages (Ref. 23), we found the posterior distributions of the dispersal parameter from its full range using Approximate Bayesian Computation (ABC)^57^ using single-hidden-layer neural networks^58^. Our mechanistic model was able to reproduce the observed alpha diversity levels (Fig. 4a), with the dispersal posteriors showing a clear increase from the Avalon to the White Sea, and a decrease in the Nama (Fig. 4, Extended Data Table 4). Our simulations demonstrate that dispersal increase is sufficient to explain much of the observed patterns of Ediacaran species richness (Fig. 4). Thus, our mechanistic model fitted by ABC seems to capture the major trends in speciation experienced through the Ediacaran suggesting that the dispersal limitation of stoloniferous reproduction is sufficient to explain the increased diversity measured between the Avalon and White Sea assemblage.

**Figure 4.**
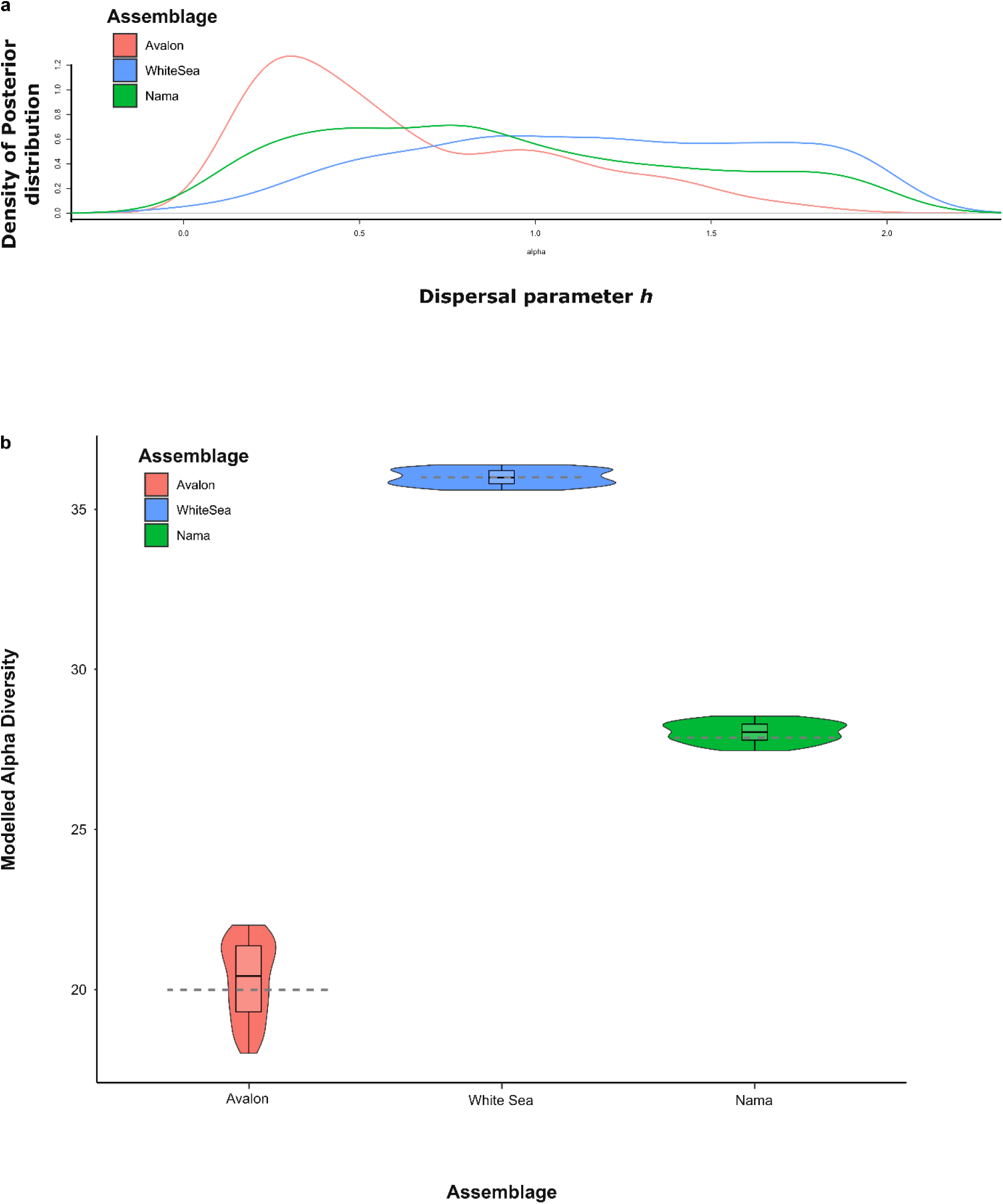
Mechanistic model simulations with rejection sampling using ABCs with neutral networks. **a)** showing the posterior distribution for the dispersal parameter for the three assemblages, with low dispersal in the Avalon, high dispersal in the White Sea and Nama assemblages. **b)** Resulting alpha diversity model simulations for the Avalon, White Sea and Nama assemblages. The grey dashed lines indicate the observed alpha diversity for the three assemblages whereby database occurrences were randomly subsampled to 50 occurrences from Ref. 23.

Our results have shown that stoloniferous reproduction in the Avalon assemblage has a significant relationship with the presence and strength of intra-specific competition. We have further demonstrated that the dispersal limitation induced by stoloniferous reproduction is the most likely underlying source of Avalonian heteromyopia, thus reducing selection pressure, with our mechanistic model indicating that these effects of stoloniferous reproduction are sufficient to explain the key trends of Ediacaran diversification. As such, stoloniferous reproduction likely played a key role in constraining early animal evolution, reducing diversification. The later onset of mobility and widespread sexual reproduction, and the consequent reduction of stoloniferous reproduction, led to an increase in competition, shifting the communities to a more niche-based dynamics.

## Acknowledgements

This work has been supported by Natural Environment Research Council Independent Research Fellowship NE/S014756/1.

## Author contributions

EGM conceived this study, which was then developed by both authors. EGM carried out the statistical analyses and wrote the first draft. Both authors carried out the model simulations. Both authors contributed to the final version of this manuscript.

## Materials & Correspondence

Figshare data will be made public on publication:

## Methods

### Quantification of isotropy of taxon populations

Isotropy (directionality of clusters) of the spatial distributions was quantified using the spatstat package^60^ in R by calculating the distribution of orientations between every point within a population and every other point^61^. These data were plotted as rose diagrams (Fig. 2 and Extended data Fig. 1), where isotropy was indicated by only small variations in the values at different angles, namely a relatively circular diagram. Strongly anisotropic distributions were highly non-circular, i.e. exhibited large variations. This variation was captured using the Inter-Quartile Range, where large values indicate high levels of anisotropy, and low values indicate isotropy. As an example of how this metric captures directionality and varies due to stoloniferous reproduction and waterborne propagation we have run our analyses on spatial data from three examples. First, an extant hydroid stoloniferous colony^62^ to demonstrate the high isotropy of stoloniferous reproduction and then two areas of an extant deep-sea coral and sponge community^63^ exhibiting high anisotropy (Extended data Figure 1), with the anisotropy varying due to different current direction on the two different areas.

### Intraspecific and interspecific species spatial distributions

The data used in this study has already been published^17,35^ and so some SPPA have already been performed on these data as follows:

‒ D and E surface: intra-specific spatial distributions up to 0.5m ^14^ and 2.0m^17^; E surface inter-specific up to 4.5m^16^.
‒ LMP inter-specific up to 1.0m^15^.
‒ Bed B: Intra and inter-specific up to 0.5m^17^.
‒ H14: intra-specific spatial distributions up to 0.5m^14^ and 2.0m^16^.
‒ Bristy Cove: intra-specific spatial distributions up to 0.25m^16^.
‒ St Shotts: intra-specific spatial distributions up to 1.5m^16^.

This published work was extended in this study by 1) extending the spatial scales of the Bed B analyses from 0.5 to 3.5m; 2) analyzing the intra-specific spatial distributions from LMP and 3) performing new intra-specific analyses on the G surface *Bradgatia* population (data Ref. ^35^). All intra and inter-specific spatial distributions were (re)analysed in order to extract the necessary data needed for the regression analyses, and to ensure consistency across the dataset.

For all 20 populations the density, maximum specimen height was recorded (Extended data Table 1). For all populations the number of normally distributed cohorts within the populations were found^64^ by fitting height-frequency distribution to various models, followed by comparison of (logarithmically scaled) Bayesian information criterion (BIC) values, which we performed in R using the package MCLUST ^64^ (Extended data Table 2). A BIC value difference of >10 corresponds to a “decisive” rejection of the hypothesis that two models are the same, whereas values <6 indicate only weakly reject similarity of the models^65^. For each population the percentage within each size class was recorded in order to indicate the number of juveniles within each population.

In order to extract the spatial scale of aggregation and/or spatial segregation, spatial analyses were performed in R using the package spatstat^60^ (described in detail in reference Ref. ^66^). The following information was extracted from the spatial distributions (Extended data Table 1) to be input into the regressions.

To determine whether spatial distributions showed significant deviations from CSR, pair correlation functions (PCF) were used as follows:

1. Determined the spatial scales and magnitudes of non-random spatial patterns by:

a. Running 999 Monte Carlo simulations for each species on a homogeneous background to generate simulation envelopes around the random (PCF = 1). 999 simulations were run (instead of 100, for example) so that the *p_d_* values (used for find the best-fit Thomas Cluster) could be measured in 0.001 increments. The highest and lowest 5% of simulations were excluded from the simulation envelope to exclude outliers^66^.
b. Where segregation (PCF<1) fell outside the Monte Carlo envelope, the minimum PCF value was recorded. So that larger values indicate more segregation^37^, the metric (1-PCF_Min_) was used.
c. Thomas Cluster models^37,66^ were fitted to the small scale PCF to determine the best-fit cluster size for the distribution. Thomas clusters are associated with reproductive processes^14,31,37,66^ so a Thomas Cluster sizes are good way to quantify the spatial scale of reproductive clusters, and have been established to be the dominant mode of aggregation for Avalonian communities^17^.

In order to compare how variables associated with reproduction different in influence on resource competition, we also calculated the LH* metric which quantifies the degree of patchiness within different systems^67^. The LH* is calculated by estimating how much nearest neighbour distances between the fossil specimens vary compared to expected random nearest-neighbour distributions (modelled using a homogeneous Poisson model)^67^, with high values indicating greater levels of heterogeneity.

### Testing for associations between the presence and strength of intraspecific competition and stoloniferous variables

The presence of intraspecific competition was indicated by an excursion below the Monte Carlo simulation and tested against the population metrics (Extended data Table 1). Model fit was compared using AIC in a stepwise approach^68^, with the best-fit model reported (Extended data Table 3). The strength of intraspecific competition (as indicated by the minimum PCF value of an excursion below the Monte Carlo simulation) was tested against the population metrics (SupplementaryTable 1).

### Mechanistic models with Approximate Bayesian Computation alpha diversity

In order to explore how dispersal limitation, immigration, niche overlap and environmental heterogeneity impacts alpha diversity, we utilised the metacommunity model of Ref. ^54^. This model is a lottery model with discrete time steps, which has been used to focus on sessile marine organisms to assess the impact of dispersal distances, selection pressures and spatial autocorrelation on biodiversity patterns. Following Ref. ^54^, we used a metacommunity of 100 patches on a grid network, seeded with a total number of species equal to 127 following Boag et al. 2016^40^. Because this is a mechanistic model which uses normalised parameters, it is not possible to parametrise it using statistical data extracted from our populations. We simulated 10,000 alpha diversities with the parameters randomly selected from the following ranges. We set a small immigration rate *n* = 0 to 0.1 to indicate the inevitable lack of full sampling that is inherent within the fossil record (cf. *n* = 0 in Ref. 54). Dispersal ranged between 0.01 (highly dispersal limited) and 2 (wide dispersal), niche overlap (*h*) between 0.01 (high niche overlap) and 10 (low niche overlap pressure). Spatial autocorrelation, used to indicate the impact of habitat heterogeneities on the taxa, followed Ref. 54 where low spatial autocorrelation corresponded to a random arrangement of the differences environmental conditions, high spatial autocorrelation minimised the differences in environmental conditions, and intermediate autocorrelation corresponded to the midpoint between the low and high spatial autocorrelations. We set the observed values of alpha diversity for the Avalon, White Sea and Nama assemblages as per Ref. 23 with α_avalon_ = 20; α_whitesesa_ = 36; α_nama_ = 28. For each of these observed values, we performed ABC to estimate the posterior distributions for dispersal, niche overlap, immigration and spatial autocorrelation, using neutral networks with a tolerance of 0.05 (i.e. using the best 500 fitting models based on the distance between simulated and observed alpha diversity).

## Extended data tables

**Extended data Table 1.**
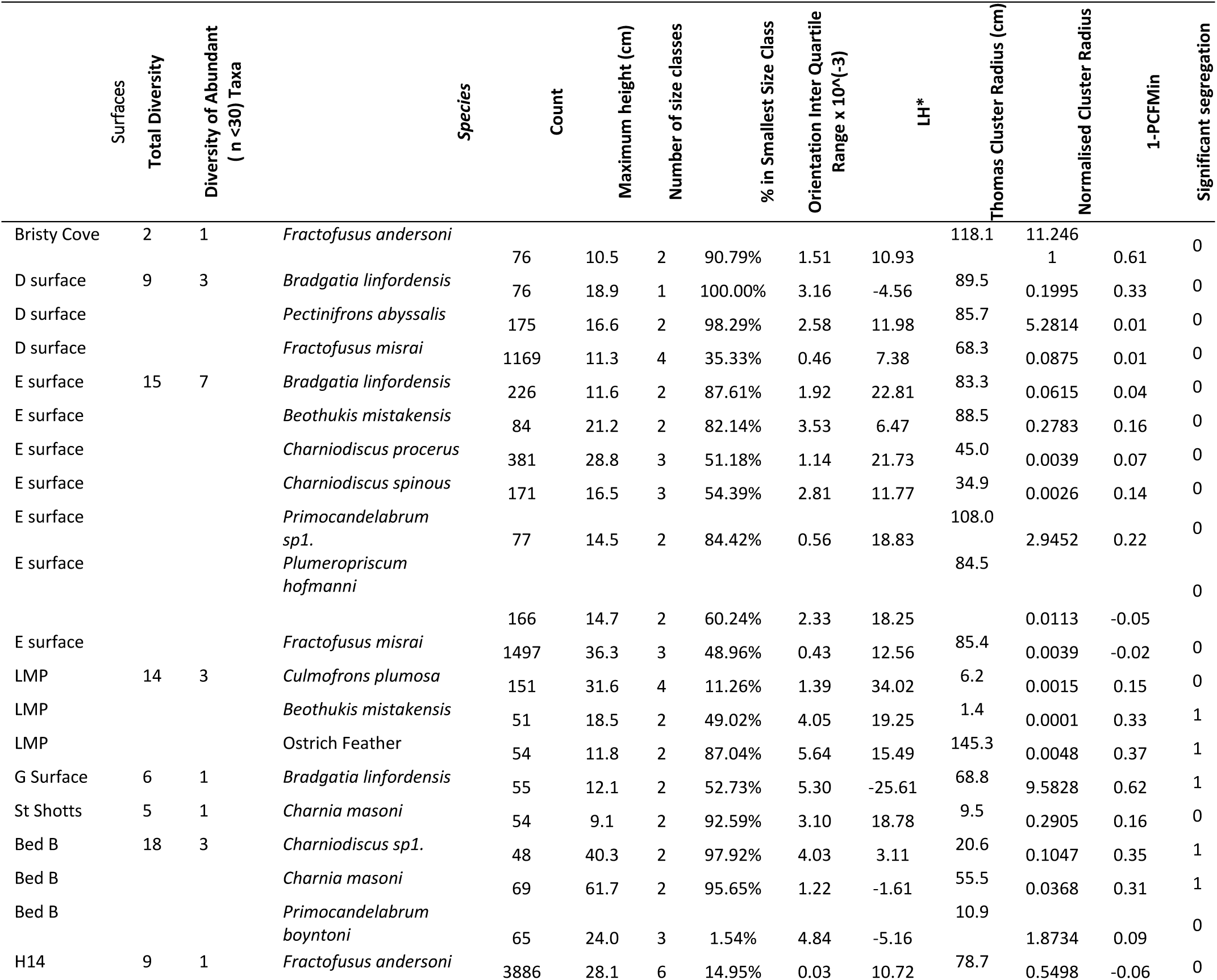
The input parameters for the binominal and linear regressions. LH* is a measure of the patchiness/heterogeneity in the data. Note that some cases, such as the G surface *Bradgatia* where Thomas Clusters are not a good model fit, the cluster size tends to zero.

**Extended data Table 2.**
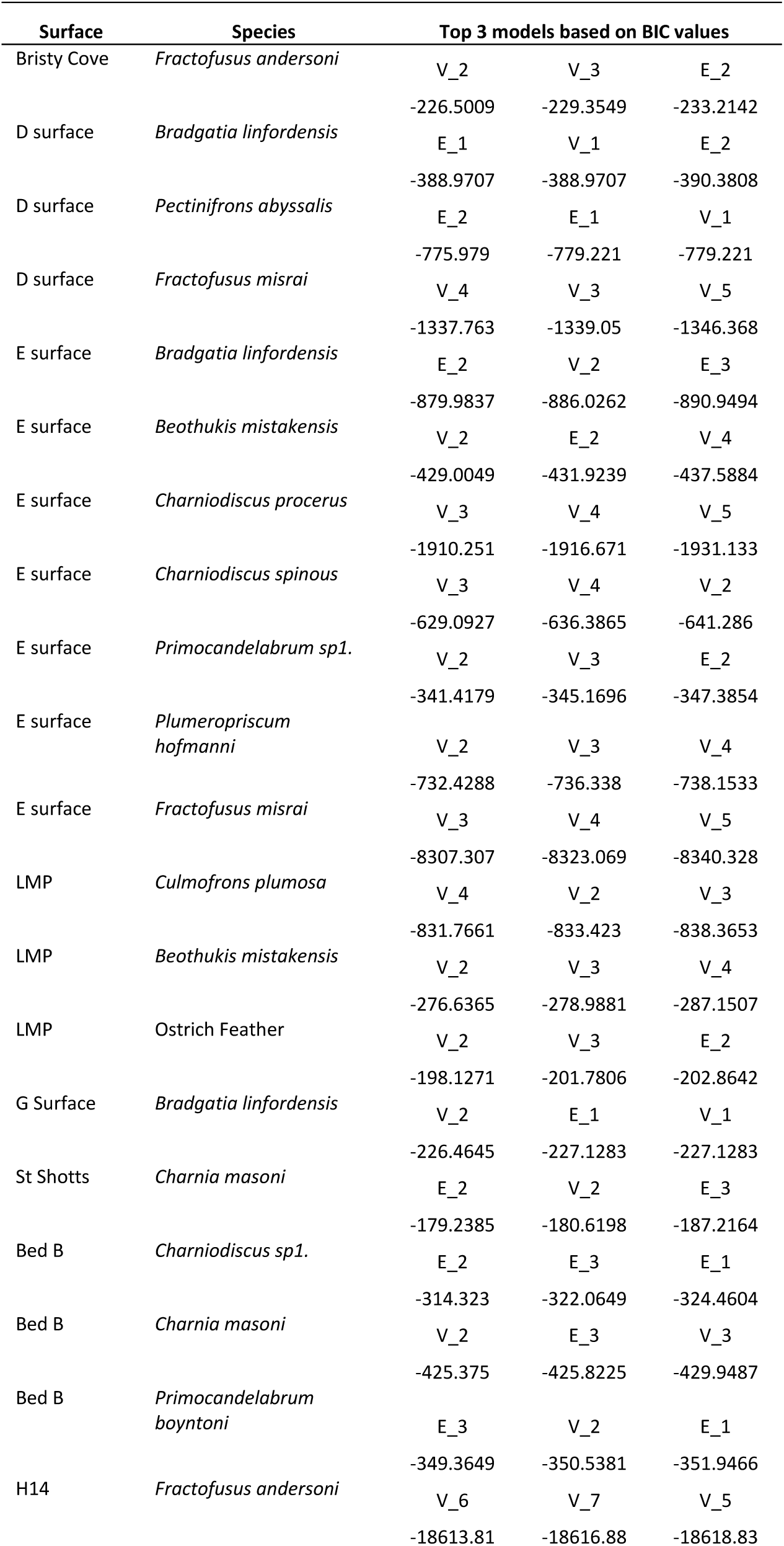
BIC values for the top 3 models found for the height distributions of the different species populations. V refers to models where the standard deviation of the best-fit model for each cohort within the population could vary, and E where the best-fit model have to have the same standard deviation. The numbers after the E or V refer to the number of cohorts in the best fit model. Note that not all the best fit models are significantly different from the second-best fit model.

**Extended data Table 3.** Summary of the stepwise regression for the variables (Extended data Table 1) showing NormCR – Normalised cluster radius; DIQR – directionality inter quartile range; PerS – percentage in the smallest size class; NSC – number of size classes and LH* – lateral heterogeneity.

| Model | *p* value | R<sup>2</sup> | AIC | ΔAIC |
| --- | --- | --- | --- | --- |
| NormCR + DIQR + PerS | 0.0021 | 0.5149 | -16.82 | 0 |
| NormCR + DIQR | 0.0014 | 0.4847 | -16.41 | 0.41 |
| NormCR + DIQR + NSC | 0.0025 | 0.5036 | -16.37 | 0.45 |
| NormCR + DIQR + PerS + NSC | 0.0064 | 0.4844 | -14.90 | 1.92 |
| NormCR + DIQR + PerS+<br>NSC+LH* | 0.0158 | 0.4542 | -13.14 | 3.68 |
| NormCR | 0.0043 | 0.3363 | -12.2 | 4.62 |
| NSC | 0.0263 | 0.2036 | -8.56 | 8.26 |
| DIQR | 0.0296 | 0.1943 | -8.32 | 8.50 |

**Extended data Table 4.**
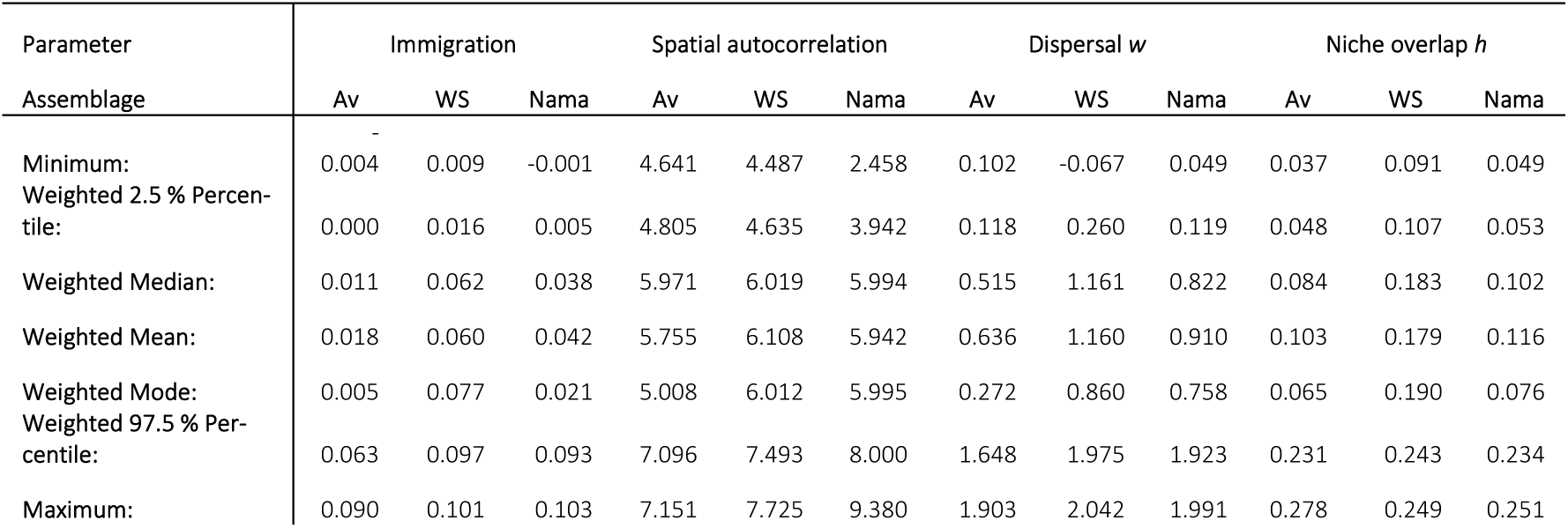
The summary statistics of the four estimated parameters of the mechanistic models.

## Extended data figures

**Extended data figure 1:**
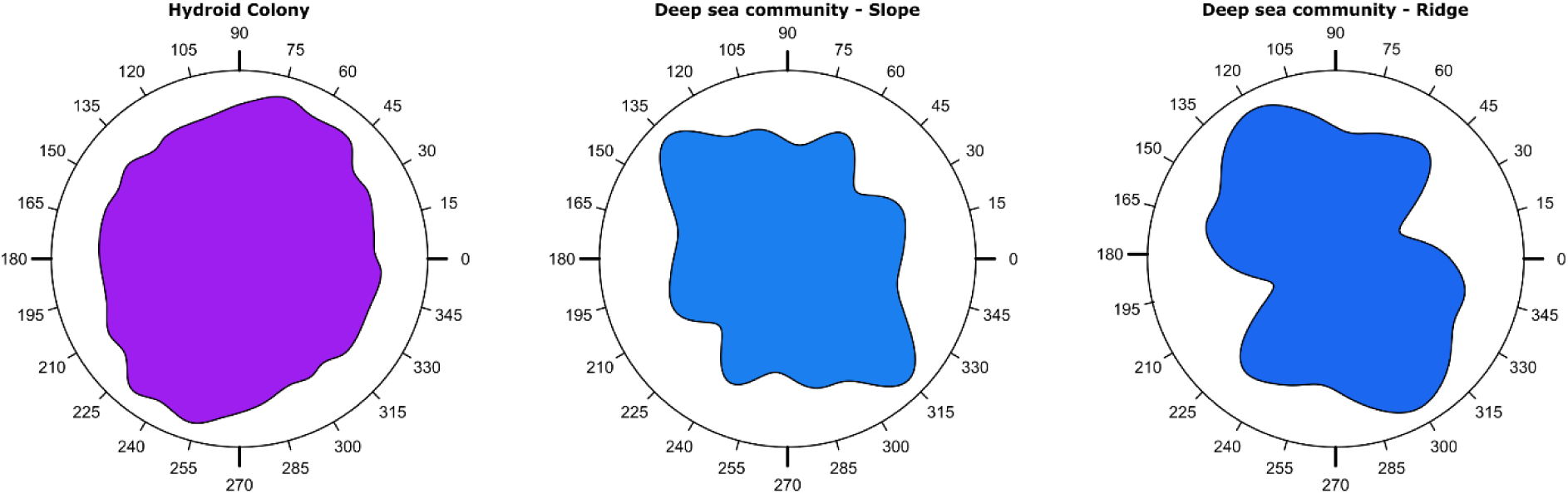
Isotropy plots for a hydroid colony connected by stolon, and two sections of a deep sea coral and sponge community subject to different current directions (from Ref. 63).

**Extended data figure 2:**
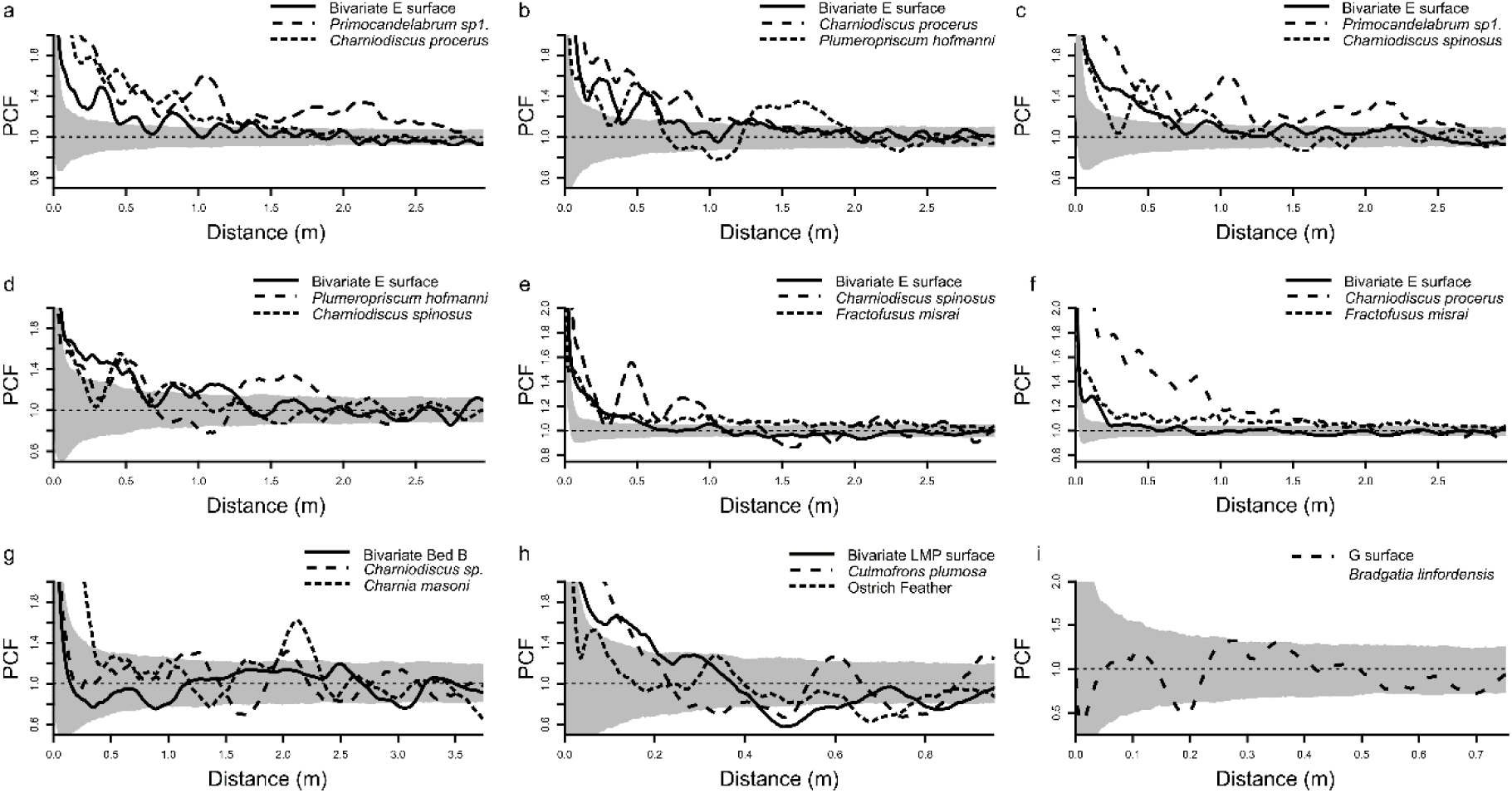
PCF for mapped taxa that exhibit intra or inter-specific segregation. For all plots the x-axis is the inter-point distance between organisms in metres. The y-axis PCF=1 indicate CSR, <1 indicates segregation, and >1 indicates aggregation. Grey shaded area depicts the bounds of 999 Monte Carlo simulations of CSR, with non-CSR patterns indicated by PCF curves are not completely within these areas. Solid lines for each plot are the bivariate PCF for the two univariate species distributions (dashed and dots respectively). Plots **a**, **b**, **f** and **g** show clear heteromyopia, where the bivariate distribution (solid line) crosses the lower bound of the Monte Carlo simulations at smaller spatial scales than the univariate distributions.

**Extended data figure 3:**
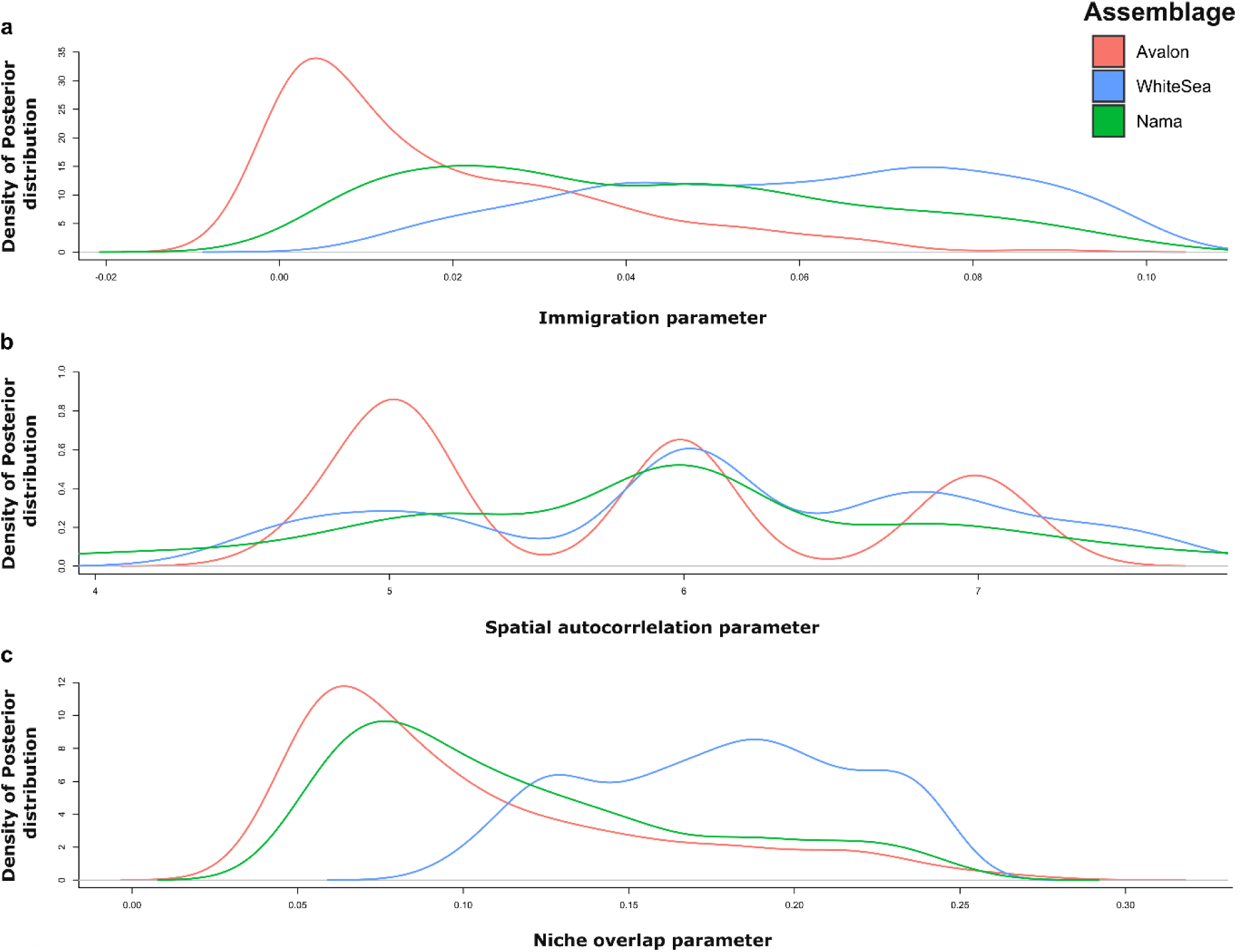
The po**s**terior distributions for the **a)** immigration parameter, **b)** spatial autocorrelation parameter and **c)** the niche overlap parameter. **a)** shows increasing immigration (although still low) between the Avalon and White Sea assemblages, with medium values for Nama. **b)** Shows the three discreet values for spatial autocorrelation (5, 6, and 7), with the Avalon showing a clear distinction of low spatial autocorrelation, with more similar values for the White Sea and Nama assemblages. **c)** shows high levels of niche overlap for the Avalon with medium to low niche overlap for the White Sea assemblage. Nama assemblage shows lower, but similar levels of niche overlap to the Avalon assemblage.

## Data availability

Data for the fossil surfaces, the modern hydroids and coral and sponge community will be made public on publication. Due to geoconservation concerns, data from Bed B cannot be made publicly available, with the full set available to researchers on request from the corresponding author. Code is available: https://github.com/egmitchell/diversityModelABCs

